# Measuring arterial pulsatility with Dynamic Inflow MAgnitude Contrast (DIMAC)

**DOI:** 10.1101/2021.01.08.425882

**Authors:** Joseph R. Whittaker, Fabrizio Fasano, Marcello Venzi, Patrick Liebig, Daniel Gallichan, Kevin Murphy

**Affiliations:** Cardiff University Brain Research Imaging Centre (CUBRIC), School of Physics and Astronomy, Cardiff University, Cardiff, CF24 4HQ, United Kingdom; Max Planck Institute for Human Cognitive and Brain Sciences, Leipzig, Germany; Siemens Healthineers, Camberly, United Kingdom; Siemens Healthcare GmbH, Erlangen, Germany; CUBRIC, School of Engineering, Cardiff, United Kingdom

**Keywords:** magnetic resonance imaging, pulsatility, cerebral arteries, echo-planar imaging, arterial stiffness, inflow effect, cerebral blood flow velocity

## Abstract

The pulsatility of blood flow through cerebral arteries is clinically important, as it is intrinsically associated with cerebrovascular health. In this study we outline a new MRI approach to measuring the real-time pulsatile flow in cerebral arteries, which is based on the inflow phenomenon associated with fast gradient-recalled-echo acquisitions. Unlike traditional phase-contrast techniques, this new method, which we dub **D**ynamic **I**nflow **Ma**gnitude **C**ontrast (DIMAC), does not require velocity-encoding gradients as sensitivity to flow velocity is derived purely from the inflow effect. We achieved this using a highly accelerated single slice EPI acquisition with a very short TR (15 ms) and a 90° flip angle, thus maximizing inflow contrast. We simulate the spoiled GRE signal in the presence of large arteries and perform a sensitivity analysis to demonstrate that in the regime of high inflow contrast it shows much greater sensitivity to flow velocity over blood volume changes. We support this theoretical prediction with *in-vivo* data collected in two separate experiments designed to demonstrate the utility of the DIMAC signal contrast. We perform a hypercapnia challenge experiment in order to experimentally modulate arterial tone within subjects, and thus modulate the arterial pulsatile flow waveform. We also perform a thigh-cuff release challenge, designed to induce a transient drop in blood pressure, and demonstrate that the continuous DIMAC signal captures the complex transient change in the pulsatile and non-pulsatile components of flow. In summary, this study proposes a new role for a well-established source of MR image contrast and demonstrates its potential for measuring both steady-state and dynamic changes in arterial tone.

**Highlights:**

- We present a novel method for measuring pulsatility of cerebral arteries.
- The inflow effect on fast GRE imaging can be exploited to yield a flow velocity dependent signal.
- We measure pulsatile flow through cerebral arteries dynamically on a beat-to-beat basis.
- We use physiological challenges to demonstrate sensitivity to dynamic and steady-state changes in vascular tone.

## 1. Introduction

Healthy functioning of the human brain depends critically upon perfusion, i.e., cerebral blood flow (CBF), which in turn is highly dependent on the health of cerebral arteries. Across the brain, CBF is tightly coupled to local metabolic demands, and can be actively regulated via arteriole smooth muscle and capillary pericyte action (Hall et al., 2014). At a more global level, cerebral autoregulation operates via a multitude of neurogenic, metabolic and mechanical factors to keep CBF across the whole brain stable in response to systemic cardiovascular stressors (Cipolla, 2009). Flow though cerebral arteries therefore varies over multiple time scales, from relatively low frequency fluctuations on the order of minutes (Zhang et al., 2000), to dynamic alterations in response to beat-to-beat blood pressure changes (Aaslid et al., 1989). Furthermore, the pulsatile pressure generated by the heart manifests as periodic fluctuations in both arterial cerebral blood volume (CBV), due to passive vessel diameter changes mediated by compliance of the vessel wall, and cerebral blood flow velocity (CBFV). Thus arterial flow has both pulsatile and non-pulsatile components, both of which are relevant in determining CBF and its regulation.

The pulsatile component of arterial blood flow, henceforth referred to as “arterial pulsatility”, is clinically relevant as it is intrinsically linked to cardiovascular health, particularly age associated loss of elasticity, i.e. arterial stiffening (Mitchell et al., 2010; Wilkinson et al., 2015). Compliant (elastic) arteries, specifically large conduit vessels like the aorta, partially buffer pulsatile energy generated by the heart, which is believed to protect the microcirculation from damage. Naturally this effect diminishes as large arteries stiffen, and this is hypothesised to be the primary route by which arterial stiffening is associated with cerebrovascular disease (Mitchell et al., 2011). Modified arterial pulsatility is therefore both directly indicative of local arterial wall stiffness, but also a passive consequence of upstream stiffness, due to the increased transmission of pulsatile energy. Thus arterial pulsatility depends predominantly on the structural properties of the arterial vasculature, whereas the non-pulsatile component of arterial flow reflects the active physiological processes that modulate vascular resistance.

Arterial flow can be measured indirectly using Transcranial Doppler (TCD) ultrasound, which is sensitive to CBFV, and has sufficient temporal resolution to resolve the full velocity spectrum in real-time, including the high frequency pulsatile component (Naqvi et al., 2013). The main disadvantage to TCD is that is limited in sensitivity to only a few large intracranial arteries where there is a suitable acoustic window in the cranium, which makes it challenging to assess downstream pulsatility. In contrast, magnetic resonance imaging (MRI) is capable of wide whole-brain field-of-view (FOV) and sensitivity to flow at multiple spatial scales, and so has the potential to remedy this particular shortcoming of TCD. Phase contrast (PC) based MRI methods quantify the phase shift of spins moving along a magnetic field gradient, and were developed in the early days of MRI (Bryant et al., 1984; Nayler et al., 1986; van Dijk, 1984), becoming the preferred approach for measuring pulsatile arterial flow. However, the need to use additional magnetic field gradients to encode velocity into the phase of the complex image data comes at the cost of acquisition speed and places a fundamental limitation of the achievable temporal resolution. The pulsatile component of flow has traditionally only been resolved using cardiac gating techniques that average data over multiple cardiac cycles (Pelc et al., 1991), yielding time averaged estimates of pulsatility insensitive to beat-to-beat variability. However, more recently PC acquisitions have been developed with sufficient temporal resolution to resolve the pulsatile components of flow in real-time, in an effort to directly image the physiologically meaningful variability that occurs over the cardiac cycle (Markl et al., 2016).

In this work, we outline a new approach to measuring arterial flow, which like real-time PC methods is simultaneously sensitive to both pulsatile and non-pulsatile components, that is based on the magnitude of the spoiled gradient-recalled-echo (GRE) MRI signal, here termed Dynamic Inflow MAgnitude Contrast (DIMAC). DIMAC exploits the inflow phenomenon, or time-of-flight (TOF) contrast, which is present during rapid GRE imaging due to the differential effect repeated RF excitation has on the degree of saturation in the longitudinal magnetisation of different spin groups according their coherent motion through the imaging plane. This saturation effect engenders sensitivity to arterial CBFV in MR image, which has traditionally been exploited to visualize vascular networks in the form of TOF angiography (Hartung et al., 2011). However, if a dynamic imaging approach is taken to collect a time series of images, this inflow contrast could theoretically be used to monitor fluctuations in arterial flow. Starting from the Bloch equations, Bianciardi et al derived analytic expressions for the spoiled GRE signal with respect to inflow (Bianciardi et al., 2016), and defined three different regimes according to whether CBFV or CBV effects dominated. In this feasibility study, we explore further this regime in which pulsatile CBFV effects are predominant and consider how this well known source of image contrast may be utilized to record arterial pulsatile flow dynamically in real-time.

We show with this approach that we can measure pulsatile arterial flow as is done with real-time PC-MRI, but with better temporal resolution, which allows us to image dynamic changes in pulsatile flow and resolve beat-to-beat pulsatile flow waveforms with high fidelity. Although we use major intracranial arteries as an in-vivo test case (namely internal carotid and middle cerebral arteries), as DIMAC is intrinsically based on saturating static tissue spins, it is theoretically less sensitive to extraluminal partial volumes than PC-MRI. This property of the DIMAC signal, which is supported by our simulation results, implies that it may show particular promise when applied to smaller arteries in the brain that are not as accessible with PC-MRI. Thus, we present a proof-of-concept for a new technique that is similar to real-time PC-MRI methods but may have particular benefits in applications focussed on dynamic flow changes and pulsatility measurement in the smaller arteries of the brain.

## 2. Methods

### 2.1. Theory

#### 2.1.1. Inflow effect

The inflow phenomenon, also called time-of-flight (TOF) effect, is fundamental to the nuclear magnetic resonance (NMR) method. Even before the advent of MRI, it had been shown that the apparent longitudinal magnetisation relaxation time T_1_ of flowing spin is shorter than stationary ones (Suryan, 1951). The mechanism behind the effect is partial saturation of the longitudinal magnetisation of stationary spins due to continual short interval RF excitation (i.e. short repetition time (TR)). In contrast, spins flowing into the imaging plane/slab do not experience this saturation effect to the same degree and so produce a stronger signal. In the extreme case of very high flow velocities (or thin slices), spins are completely refreshed between RF pulses, and so the fully relaxed equilibrium magnetisation is available to be measured in the transverse plane. In the more general case, assuming transverse magnetisation is spoiled after readout, different spin isochromats reach different steady states of longitudinal magnetisation determined by their flow velocity.

The literature already includes detailed quantitative analysis of the short TR spoiled GRE MRI signal steady state (Brown et al., 2014), including the effect of flow velocity in non-static spins (Bianciardi et al., 2016; Gao et al., 1988). In the simplest case in which a slice of thickness L is orientated perpendicularly to a blood vessel, which can be modelled as a cylinder, assuming plug flow and a 90° flip angle, the longitudinal magnetisation scales linearly with velocity v and is given by

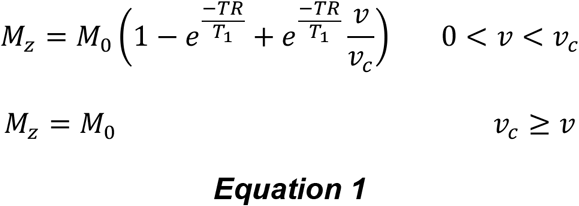

where M_0_ is the equilibrium magnetisation, and v_c_ is the critical velocity, above which the flowing spins experience only one RF pulse when crossing the excited slice. The measured signal is simply scaled by the spin density/volume and a transverse relaxation factor. The critical velocity v_c_ is determined by the ratio of slice thickness and TR (L/TR). If v > v_c_ there is no longer flow dependence, and the longitudinal magnetization remains at equilibrium, in the steady state. Based on this theory, if v < v_c_ we hypothesise that fast spoiled GRE sequences may prove very useful for measuring pulsatile flow in arteries with high temporal resolution.

#### 2.1.2. Sensitivity analysis - simulations

The cardiac induced pressure waveform that propagates through the vasculature consequently leads to pulsatile flow in arteries (Wagshul et al., 2011), which manifests as pulsatile changes in both CBV and CBFV. In this section, we perform simulations to assess the sensitivity of the spoiled GRE signal to pulsatile changes in CBFV and CBV. The cardiac phase (*τ*) dependent signal can be modelled in a two-compartment model simply as the sum of intraluminal (*i*) and extraluminal (*e*) compartments

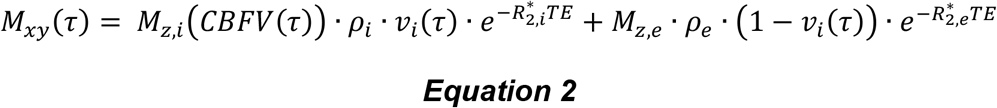

where *M_z_* is the longitudinal magnetization defined in Equation 1 (which is a function of CBFV for intraluminal spins), *ρ* is the spin density for the respective compartments, and *v_i_* is the intraluminal volume (i.e. CBV). Cardiac pulsatile physiology dictates that both CBV and CBFV are functions of *τ*, and so too is the measured signal. The maxima of CBFV and CBV occur during the systolic peak (*sys*) of the cardiac phase, with baseline values observed during the diastolic (*dia*) portion. If during baseline a voxel is entirely contained with the vessel lumen (i.e. *v_i_* (*τ_dia_*) = 1), then the measured signal will be sensitive only to changes in CBFV over the cardiac cycle. Thus, the signal is only sensitive to pulsatile CBV in voxels with an extraluminal partial volume (i.e. *v_i_* (*τ_dia_*) < 1), and the degree to which it is depends on both the baseline partial volume and the maximum partial volume (i.e. systolic peak).

We simulate the conditions for a single voxel (with dimensions 2×2×10 mm) orientated perpendicularly to the middle cerebral artery (MCA), which we model as a straight cylinder. The extraluminal partial volume (*v_e_*) is assumed to be composed entirely of CSF, which is the most appropriate assumption for large arteries in the brain (Dieleman et al., 2014). All MR simulation parameters are listed in Table 1. MCA CBFV varies both with cardiac cycle (i.e. pulsatility), but also across the vessel lumen due to the laminar flow profile, being strictly 0 at the vessel wall, and peaking during systole at the centre. Thus, CBFV was allowed to vary in the range between 0 – 100 cm s^-1^ to provide a realistic distribution of flow velocities (O’Rourke et al., 2020). Measuring *in-vivo* diameter changes within intracranial arteries is challenging, but high-resolution images obtained with ultra-high field MR provide the best estimates. Using this technique, it has been estimated that the MCA changes in cross sectional area by ~2.5% over the cardiac cycle (Warnert et al., 2016). Assuming a perfect cylinder this translates to a change in CBV of the same magnitude. For simulations, we assumed that CBV could increase by up to 5% of its baseline *diastolic* value. Using the above physiological ranges, we assess the global sensitivity of the DIMAC signal to changes in CBFV and CBV as follows;

1. We define baseline CBFV and CBV values and calculate the signal magnitude (S).
2. We then randomly sample (1000 samples) ΔCBFV and ΔCBV values uniformly from the physiological plausible range and calculate the change in signal from baseline (ΔS).
3. We then regress ΔS against ΔCBFV and ΔCBV (normalized between 0 and 1) to estimate regression coefficients that are in units of ΔS per dynamic range of CBFV and CBV.
4. The ratio of the regression coefficients (ΔS/ΔCBFV divided by ΔS/ΔCBV) expresses the relative sensitivity to pulsatile changes in CBFV over CBV.

**Table 1:** MR parameters for simulation at 3T.

| Parameter | Value | Reference | Comment |
| --- | --- | --- | --- |
| Arterial blood $T_1$ | 1664 ms | (Lu et al., 2004) | At Hct = 0.42 |
| Arterial blood $T_2^*$ | 48.4 ms | (Zhao et al., 2007) | At $sO_2=98\%$ and Hct = 0.44 |
| CSF $T_1$ | 3817 ms | (Lu et al., 2005) | |
| CSF $T_2^*$ | 400 ms | (Pinto et al., 2020) | |
| Arterial blood $\rho$ | 0.85 | (Herscovitch and Raichle, 1985) | Assume Hct = 0.44 |
| CSF $\rho$ | 1 | | Assume like water |

#### 2.1.3. Impact of SNR

We simulated different levels of SNR to explore how the predictions of the sensitivity analysis would translate to the *in-vivo* scenario. The beat-to-beat pulsatile CBFV is modelled with a Fourier basis set as

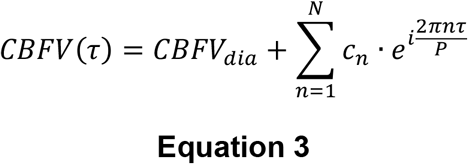

where *CBFV_dia_* is the baseline (i.e. dc component) during diastole, *P* is the beat-to-beat time period, and *n* is the harmonic of the fundamental frequency. A total of *N* = 10 harmonics were used taken from (Yang et al., 2019) in order to model a generic pulsatile flow response.

### 2.2. Experimental Protocol – in-vivo imaging

We performed two separate *in-vivo* MRI experiments using a highly accelerated GRE echo-planar-imaging (EPI) acquisition to demonstrate the potential of the DIMAC method as follows: i) an experiment to estimate the steady-state cardiac induced pulsatile flow response in the middle cerebral artery (MCA), using a hypercapnia challenge to demonstrate the sensitivity of DIMAC to subtle changes in vascular tone (*HC-challenge* experiment); ii) an experiment to measure flow changes in the Internal Carotid (ICA) and Vertebral (VA) arteries during a thigh cuff release (TCR) challenge to demonstrate sensitivity to dynamic changes in the non-pulsatile component of CBFV, and beat-to-beat pulsatile flow (*TCR-challenge* experiment). Finally, we also performed a simple flow phantom experiment with the same acquisition in order to verify the strong flow velocity dependent signal that is predicted in the very short TR domain (included in *Supplementary material*).

#### 2.2.2. Imaging protocol

All experiments described below were performed on a Siemens 3T MAGNETOM Prisma clinical scanner with a 32-channel receiver head-coil (Siemens Healthcare GmbH, Erlangen), and used a prototype single slice GRE EPI sequence, with the number of repetitions varied according to the experimental requirements (see following sections). The protocol was optimized for maximum sensitivity to the inflow effect by making the TR as short as possible, which included removing fat saturation pulses. Acquisition parameters were as follows: flip angle=90°, FOV=192mm (2 mm^2^ in-plane resolution), GRAPPA=5, partial Fourier = 6/8, TR=15ms, TE=6.8ms, slice thickness=10mm. For all in-vivo experiments standard TOF scans were performed in order to guide the placement of DIMAC slices perpendicularly to the artery of interest. All participants gave written informed consent, and the School of Psychology Cardiff University Ethics Committee approved the study in accordance with the guidelines stated in the Cardiff University Research Framework (version 4.0, 2010). Data are publically available through the Open Science Framework (DOI 10.17605/OSF.IO/ZQ5E3).

#### 2.2.3. In-vivo experiments

##### 2.2.3.1. HC-challenge experiment

An experiment was performed in 5 healthy participants to demonstrate the sensitivity of DIMAC to measuring arterial pulsatility in-vivo, with slices positioned to target the MCA as shown in Fig.1A. A hypercapnia challenge (HC) in which two distinct levels of partial pressure of end-tidal CO_2_ (P_ET_CO_2_) were targeted was used as a global vasodilatory stimulus in order to model changes in vascular tone.

**Figure 1:**
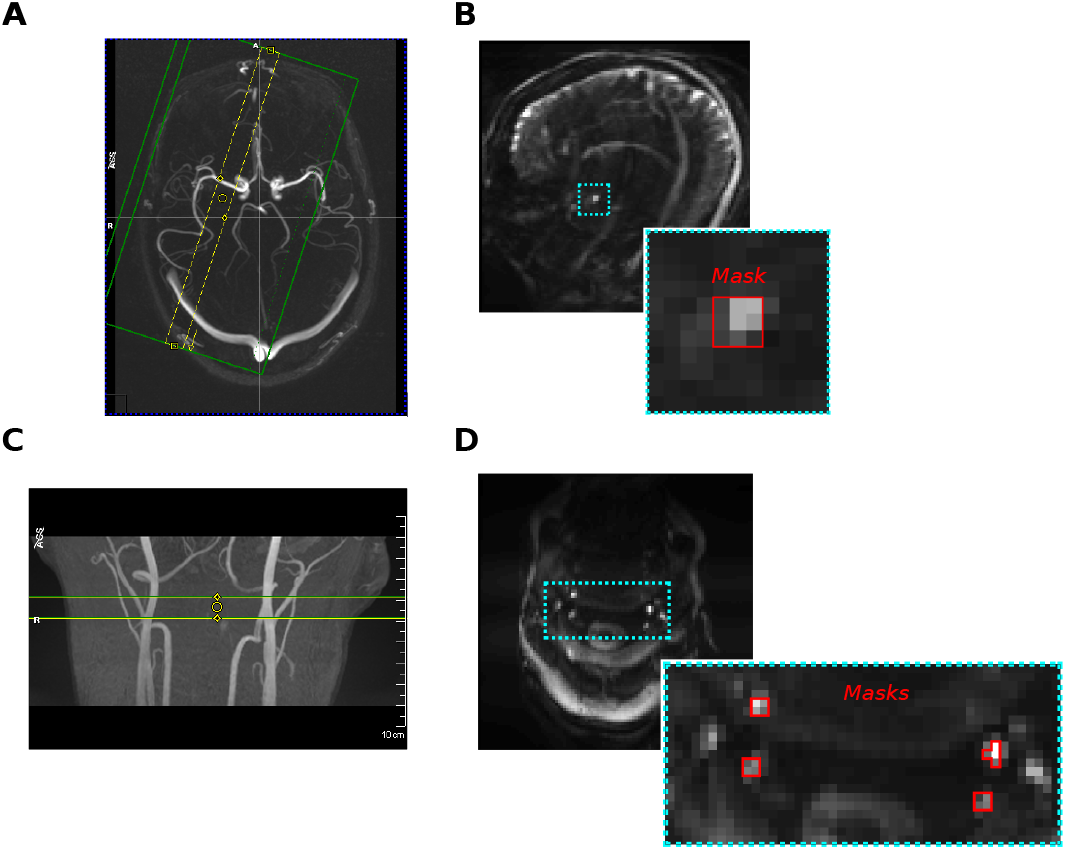
A) An example of the slice placement at the M1 segment of the MCA for the *HC-challenge* experiment. B) An example DIMAC image, including MCA mask. C) Slice placement for the subject in the *TCR-challenge* experiment placed to include best perpendicular placement of bilateral ICAs and VAs. D) Subject’s DIMAC EPI image and artery masks.

For each subject a series of scans was performed, which included acquisitions with the default protocol (TR = 15 ms, 4096 repetitions), and 3 modified acquisitions in which the TR was increased by a factor of 2 (i.e. TR = 30,60, and 120 ms). These were included to provide empirical support for the theoretical prediction that sensitivity to arterial pulsatility will decrease at longer TRs. The number of repetitions for these modified acquisitions was also adjusted by a factor of 2 (i.e. 2048, 1024, and 596 repetitions), such that the scan length was always exactly 61.44 seconds. In addition to modifying TR, scans were also performed at 3 distinct levels of P_ET_CO_2_, in which levels were defined with respect to individual subject baseline, which was determined during the initial set up period during the experiment. For each TR protocol, acquisitions were repeated at normocapnia (+0 mm Hg w.r.t. baseline), and two levels of hypercapnia (+4 and +8 mm Hg w.r.t. baseline). Thus, for each subject a series of twelve scans in total was performed, i.e. each of the 4 TR protocols at each of the 3 levels of hypercapnia. The experiment therefore had a factorial design with 2 factors (TR and HC) with 4 and 3 levels respectively (TR15, TR30, TR60, and TR120; HC0, HC4, and HC8). For each HC level the order of scans was the same (TR15, TR30, TR60, TR120), but the order of HC levels was randomised across subjects.

The details of the HC experiment are as follows. A tight fitting mask was used to manually deliver gas through a system of custom-made flow meters, as previously described (Whittaker et al., 2016). A sampling line connected to the mask was used to monitor P_ET_CO_2_ levels, and flow of medical air and 5% CO_2_ was manually adjusted to target discrete levels (+4 and +8 mm Hg) above the participant’s predetermined baseline value. The baseline level was determined on an individual subject basis at the beginning of each scanning session. The mask circuit setup allowed gases to mix in a length of tubing before reaching the mask, and a minimum total flow rate of 30 L/min was maintained at all times. For the normocapnia scans only medical air was delivered. For hypercapnia scans the flow rates were adjusted to achieve the desired target prior to the start of the acquisition, with sufficient time given to ensure a steady state was reached. Flow was always returned to medical air in between HC levels to allow subjects to return to baseline, and subjects were given ~1-2 min of recovery at baseline between hypercapnic levels. The start of a new hypercapnic level and delivery of CO_2_ gas was always preceded with the subject’s verbal consent. For each hypercapnic period, at least 1 minute was allowed when transitioning to a new P_ET_CO_2_ level in order to ensure a steady state at the target end-tidal value. As there were different TR protocols for each HC level, each lasting ~1 min, when factoring in transitions, subjects were only ever at a particular hypercapnic level for ~5-10 min. S2 in the *Supplementary material* shows an example P_ET_CO_2_ trace and relative scan timings for one subject. All subjects tolerated the HC challenge well and none reported any significant discomfort. Additionally, photoplethysmography (PPG) traces were recorded concurrently to provide an independent measure of the cardiac cycle.

##### 2.2.3.2. TCR-challenge experiment

An experiment was performed in a single subject to demonstrate the utility of DIMAC for measuring changes in flow/pulsatility dynamically. In order to modulate flow we used a thigh cuff release (TCR) challenge, as it is known to cause a robust transient drop in blood pressure (Aaslid et al., 1989; Mahony et al., 2000). A single transverse slice was placed in the neck at a position approximately perpendicular to both the internal carotid arteries (ICA) and vertebral arteries (VA), as shown in Fig. 1C. The TCR protocol, detailed here in conference abstract form (Whittaker et al., 2020), was briefly as follows: Pneumatic cuffs were placed around the tops of both thighs and inflated to +40 mm Hg above baseline systolic BP pressure for 152 s and then rapidly deflated. Scanning of the DIMAC acquisition was timed such that data collection began 20 s before deflation, and each scan lasted ~60s (4096 repetitions). A series of 5 TCR manoeuvres were repeated, and both concurrent PPG and beat-to-beat blood pressure (Caretaker, Biopac) traces were recorded.

### 2.3. Analysis

#### 2.3.2. In-vivo experiments

Data for both in-vivo experiments were processed using AFNI (Cox, 1996) and MATLAB. All images were first motion corrected using AFNI’s 2dlmReg function and filtered to remove linear drifts. Subsequent analysis of the data was performed on ROI average time series and is described below. For the TR15 condition, pulsatile CBFV weighting in the signal is sufficiently high such that the periodic signal is clearly visible and we could perform peak detection to identify each cardiac cycle without the need for the external PPG, which we refer to as *Beat-to-beat fit* and describe below. For the other TR conditions this peak detection is no longer possible, but the pulsatile component of the signal can still be extracted by using the PPG as an external reference of the cardiac phase, which we refer to as *Cardiac binned average*, and describe below. However, with this method, only a time average pulsatile component can be extracted.

##### 2.3.2.1. Cardiac binned average

As has been previously done with PC and functional MRI (fMRI) approaches, we can use the PPG as an external reference to determine in which phase of the cardiac cycle a particular MR image was acquired. *Systolic* peaks were detected in the PPG signal, and then each beat-to-beat interval was split into a set of *n* cardiac bins. Thus, individual data points in the time series could be sorted into one of these *n* bins, and then averaged into a new *n* point cardiac phase time series, representing the time averaged pulsatile component of the signal. As the sampling rate is different for each TR protocol, the number of bins *n* was varied to be approximately half the sampling rate, according to equation 4.

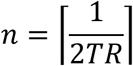

where [*x*] is the ceiling function that rounds *x* up to the next integer value. With this method we also calculated a time averaged pulsatility index (PI), a simple metric for quantifying the degree of pulsatility in the signal, and defined as the maximum value of the average signal minus the minimum value divided by the mean.

##### 2.3.2.2. Beat-to-beat fit

As is customary with pulsatile flow waveforms, a Fourier series basis set was used to model the pulsatile component of the signal. In order the detect this on an individual beat-to-beat basis the *diastolic* peak of each cardiac cycle was detected (see Fig. 2B), and then, after removal of linear trends, linear regression was used to fit Fourier basis set to the signal between each pair of *diastolic* peaks. Fig. 2A shows the average R^2^ of individual beat-to-beat fits for the *HC-challenge* experiment (averaged across HC condition and subject for the TR15 condition only). As expected the R^2^ increases with the number of harmonics included in the Fourier basis set and begins to plateau at ~6. Thus, 5 terms (the fundamental frequency + 4 higher order harmonics) was chosen as the best balance between goodness-of-fit and parsimony, as increasing to a greater number of terms offers only marginal increases in R^2^, but at the risk of over-fitting. As each individual beat is characterised by its own set of Fourier coefficients, the time averaged pulsatile waveform can be modelled by simply averaging these together, and then estimated with an arbitrary number of data points.

**Figure 2:**
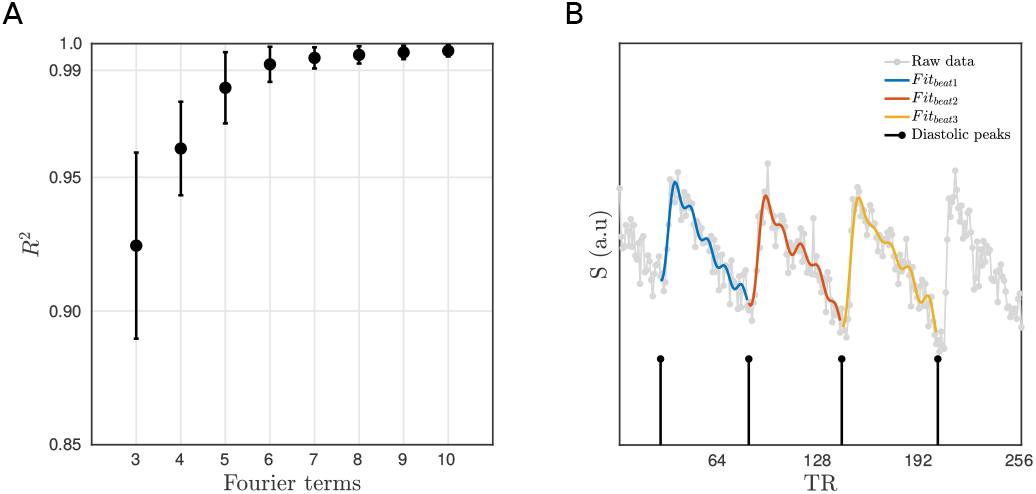
A) The average amount of variance explained in the beat-to-beat fit as a function of the number of Fourier terms used. B) An example showing detected *diastolic* peaks and the beat-to-beat pulsatile fits.

Additionally, *systolic* peaks (i.e. maximum signal during systole) were also detected. Both *diastolic* and *systolic* peaks were up-sampled to the original sampling frequency to create *diastolic* and *systolic* time series respectively. Thus, the whole time series could be deconstructed into a non-pulsatile component (i.e. the low frequency fluctuating *diastolic* peaks), and individual beat-to-beat pulsatile components that could be characterised by a set of Fourier coefficients. The *diastolic* and *systolic* time series are equivalent to the envelop of the dynamic pulsatile signal and contain low frequency information related to the physiological factors that affect the non-pulsatile component of CBFV.

##### 2.3.2.3. HC-challenge experiment

For each subject, ROIs located at the MCA were defined from the average image across all scans, as a 9 voxel mask encompassing the artery, which was selected manually such that the centre voxel was the brightest in the region of the artery (see Fig.1B). For the default protocol (TR15), each ROI time series was processed using the *Beat-to-beat fit* method described above. Fourier coefficients for each individual beat were averaged together and then the time-averaged waveform was reconstructed with 100 data points. Additionally, the ROI time series were also processed using the *Cardiac binned* average method in order to calculate and compare the PI across all TR conditions. The PI was calculated for each TR and HC condition, and then a repeated-measures ANOVA was used to test for an effect of TR on PI, after averaging across HC levels.

##### 2.3.2.2. TCR-challenge experiment

For each subject 4 ROIs were created encompassing the ICAs and VAs bilaterally as follows; the brightest voxel in the region of each artery was used to define the centre of a 5×5 voxel search space, within which the 4 brightest voxels were selected to form a mask. As shown in Fig.1.D, voxels in all masks were contiguous to create a single ROI for each artery. Average time-series were extracted from each ROI and then processed using the *Beat-to-beat fit* method described above.

## 3. Results

### 3.1. Simulations

Fig.3A shows the relative sensitivity of the GRE MRI signal as a function of TR for different flip angles. The sensitivity to CBFV increases rapidly as TR is decreased, and this is most pronounced for the maximum α=90°. Thus, we define the “DIMAC regime” as this region of the GRE parameter space that engenders high sensitivity to pulsatile CBFV. Fig.3B shows the simulated signal plotted as a function of CBFV in the case of α=90° and TR=15 ms, where it can be clearly seen that the effect of pulsatile CBV is very small compared to CBFV when v < vc, which in this simulation is 66cm/s. When v > vc, all flow sensitivity is lost and the signal is purely sensitive to changes in CBV, although at a much reduced dynamic range. It is also evident that the two parameters are coupled so that the dynamic range of CBV signal variance scales with CBFV, and that the magnitude of the signal is dependent on baseline partial volume. These results suggest that in the DIMAC regime of high saturation (TR=15ms, flip=90°), partial volume of arterial blood merely scales the overall signal magnitude, which is always relatively more sensitive to pulsatile CBFV over CBV.

**Figure 3:**
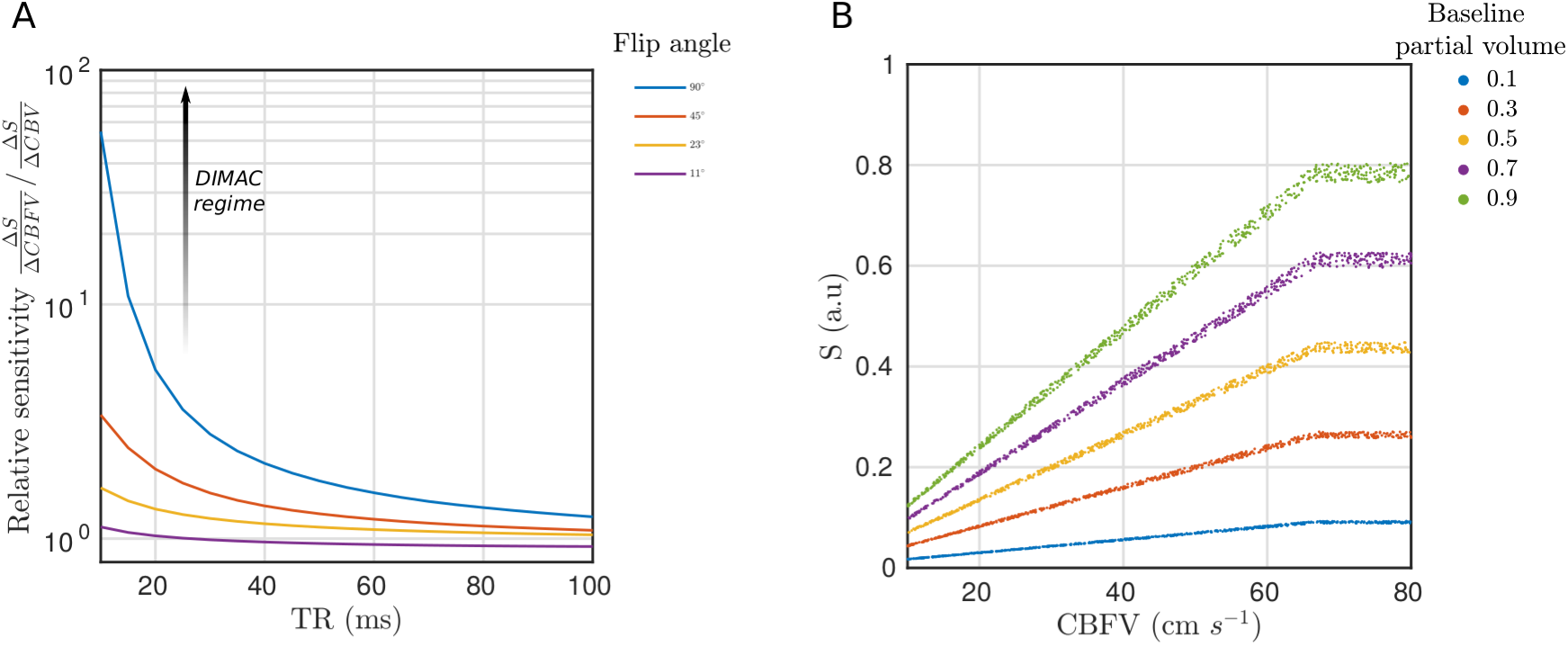
A) Relative sensitivity of spoiled GRE signal to pulsatile CBFV compared with pulsatile CBV, where each line represents a different flip angle. The annotated arrow represents how the GRE signal moves into the DIMAC regime of high sensitivity to pulsatile CBFV over pulsatile CBV as TR decreases and flip angle increases. B) Scatterplot of simulated spoiled GRE signal in the DIMAC regime of low TR (15 ms) and high flip angle (90°) as a function of CBFV. Note that the spread of data points reflects signal variance due to pulsatile CBV, which is small compared with variance due to CBFV within the normal physiological range. In every case the signal plateaus at the critical velocity 66 cm s^-1^, i.e. when flow velocity increases to the point when spins only experience a single RF pulse and the signal becomes sensitive to CBV alone.

The pulsatile signal was then modelled for different baseline partial volumes and SNR levels. As seen in Fig.4A the quality of the CBFV weighted pulsatile signal is a function of both baseline partial volume and SNR, which determines the fidelity with which single beats can be resolved. Fig.4B plots the agreement of the simulated signal with the pulsatile CBFV waveform, and it can be seen that even in the lowest partial volume and SNR case, although individual beats can’t be seen, with sufficient averaging (100 beats) the MR signal still shares more than 50% variance with the pulsatile CBFV. In cases of high SNR and baseline partial volume, easily achievable *in-vivo*, the individual beats can be resolved with high fidelity.

**Figure 4:**
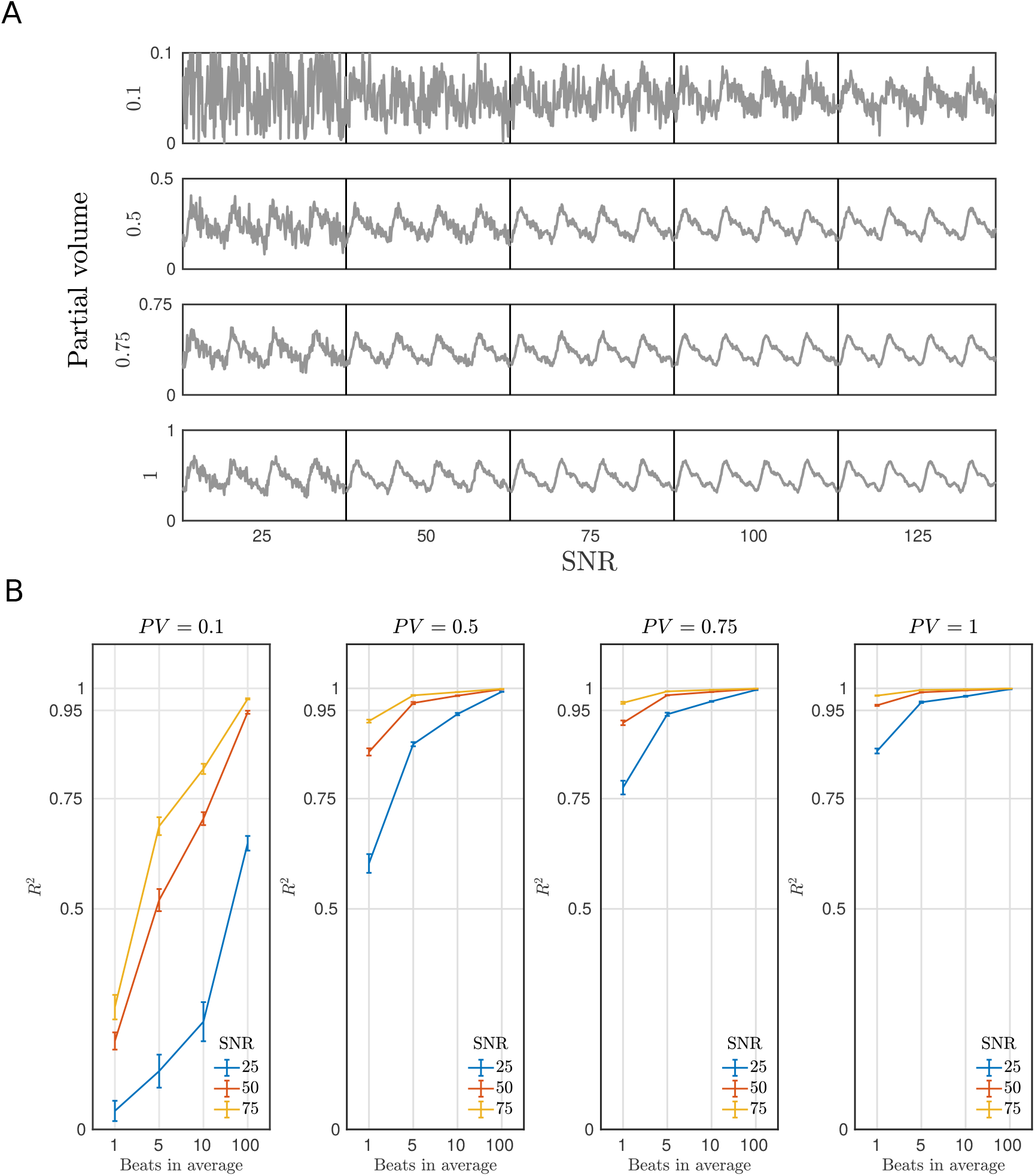
A) Simulated DIMAC signal over 4 beats. Each panel represents a different partial volume (row) and SNR condition (column). Note the different y-axis scale for each partial volume. Thus, for a given SNR, increased partial volume of arterial blood increases signal magnitude and yields a stronger pulsatile signal. B) Each panel shows the amount of variance explained in the true CBFV waveform by the simulated DIMAC signal as a function of number of beats included in the average. Naturally, as the number of beats included in the average increases more noise is averaged out, and thus the R2 increases. However this also shows that when both partial volume and SNR are sufficiently high, individual beats can be resolved with high fidelity (i.e. a high R2 with respect to the true CBFV waveform).

### 3.2. In-vivo experiments

#### 3.2.2. HC-challenge experiment

The theory predicts that in the “DIMAC regime” there is a strong sensitivity to pulsatile CBFV, and comparing the signal from the different TR conditions empirically supports this. A strongly periodic pulsatile signal is evident in the time series for the TR15 condition, but is far less visible for the TR30 condition, and not readily visible for the TR60 and TR120 conditions (see Fig 5A), as predicted by the theory. This is also reflected in the power spectra, with the power of the fundamental cardiac frequency clearly reducing as a function of TR, and higher order harmonics becoming less well defined also (see Fig 5A). Fig 5B shows the PI for each TR and HC condition. It is clear, as expected, that the PI is strongly dependent on TR condition, and this is quantitatively verified by the results of a repeated measures ANOVA on PI, which shows a significant effect of TR (p=0.00062).

**Figure 5:**
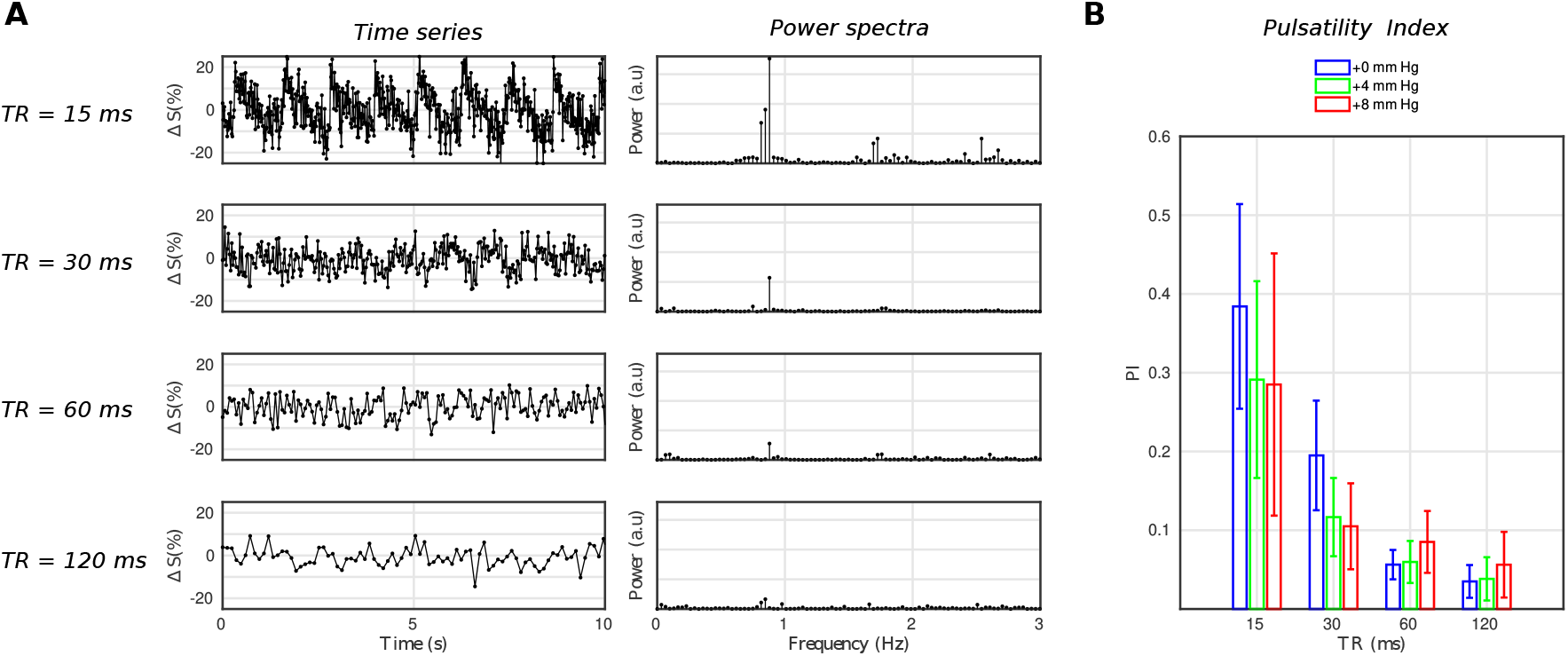
A) Example (subject 1) time series and their power spectra for each TR condition. Both the time series and power spectrum make it clear that there is a strong degree of pulsatility for the TR15 conditions, which falls away with higher TR values as predicted by the theory. B) The pulsatility index for each TR and HC condition, which quantitatively confirms that which is observed in A.

Figure 6A shows the first 15 s of the DIMAC time series for each subject in the TR15 and HC0 condition, along with the beat-to-beat fit. Qualitatively one can observe differences in the pulsatile signal shape across subjects, which is more clearly seen in the average responses for each subject in Fig. 6B. One can also see in Fig. 6A that there is beat-to-beat variability in the pulsatile signal, for example in the subject presented in the fifth row, one can see that the first few beats are qualitatively different in shape to the later beats.

**Figure 6:**
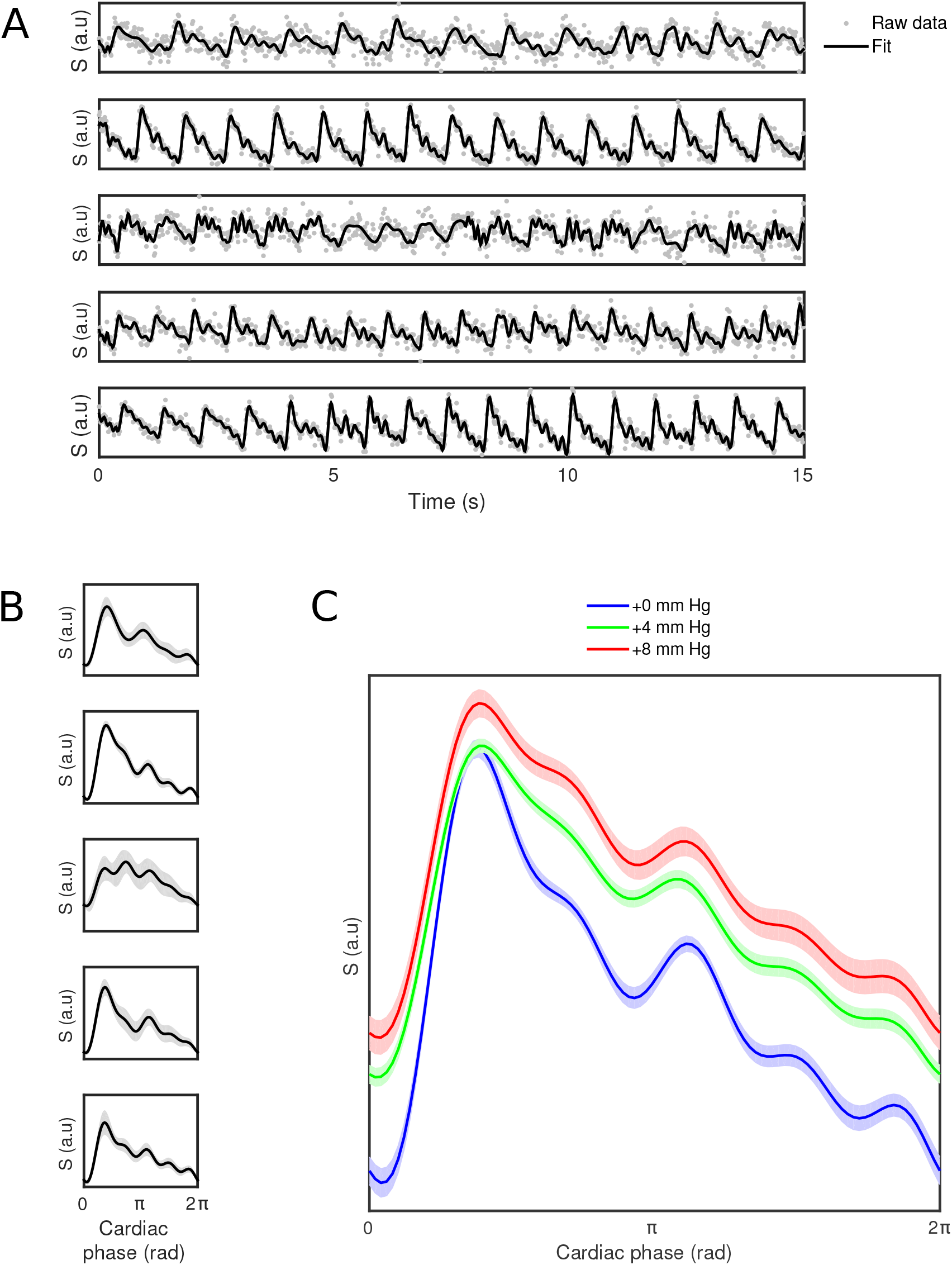
A) First 15s of time series for each subject for HC-challenge data (TR15 and HC0 condition), with beat-to-beat fit overlaid. B) The mean pulsatile waveform (±SD shaded area) for each subject (TR15 and HC0 condition). C) The across subject mean pulsatile waveforms ((±within subject error shaded area) for each HC condition of the HC-challenge. Qualitatively one can see the pulsatile waveform shape is modulated by hypercapnia.

Fig. 6C shows the group average cardiac phase waveforms across different HC conditions. The cardiac phase waveforms show at least two clear peaks, which are consistent with what is observed with TCD (Kurji et al., 2006; Robertson et al., 2008), with the first one representing the systolic peak, and the second one representing a reflection wave, preceded by the so called “Dicrotic notch”, related to a transient increase in pressure associated with the aortic valve closing. Qualitatively, there is a clear modulation of the waveform baseline with increasing levels of hypercapnia, due to increased flow velocity, and also clear modulation of the waveform shape. With increased hypercapnia the two peaks become broader, and less clearly separated from one another. There is also evidence of an additional reflection peak appearing on the downward slope of the primary peak, which becomes more pronounced with increasing hypercapnia and has previously been demonstrated with TCD (Robertson et al., 2008). Closer peaks may indicate resonance effects produced by pressure waves in shorter vascular systems (Nichols et al., 2011).

#### 3.2.2. TCR-challenge experiment

Fig. 7A shows the first 15 seconds of the L-ICA DIMAC signal along with the beat-to-beat fit and the signal *diastolic* and *systolic* peaks. A very strong periodic signal is clearly observable, with the prominent systolic and wave reflection peaks discernable on a beat-to-beat basis. Figure 7B shows the full *systolic* and *diastolic* peak time courses of the signal during the thigh cuff release challenge in both Ieft and right ICA and VAs. The magnitude of the signal changes are larger in the ICAs than in the VAs, and the traces appear smoother, which is perhaps not surprising given that the ICAs are larger arteries (diameter of ~5 mm compared with ~3 mm for VA), and thus will have higher SNR. The event-locked change in the CBFV evoked by the TCR is present in the signal *systolic* and *diastolic* peak time courses, and is clearly seen bilaterally in the ICA, but only partially in the VA, primarily in the left branch. Interestingly, it is also clear that the TCR response in the ICA *systolic* peak time series is delayed with respect to the *diastolic* peak time series. Furthermore, the *diastolic* peak response shows a marked drop in amplitude, followed by a subsequent overshoot, whereas the *systolic* peak response shows a simple slowly evolving increase in amplitude and return to baseline. *Supplementary material* S3 examines the TCR response more closely in the ICA DIMAC *systolic* and *diastolic* peak time series and compares them with the heart rate and mean arterial blood pressure responses, which demonstrates that these DIMAC signal changes are clearly of a physiological origin.

**Figure 7:**
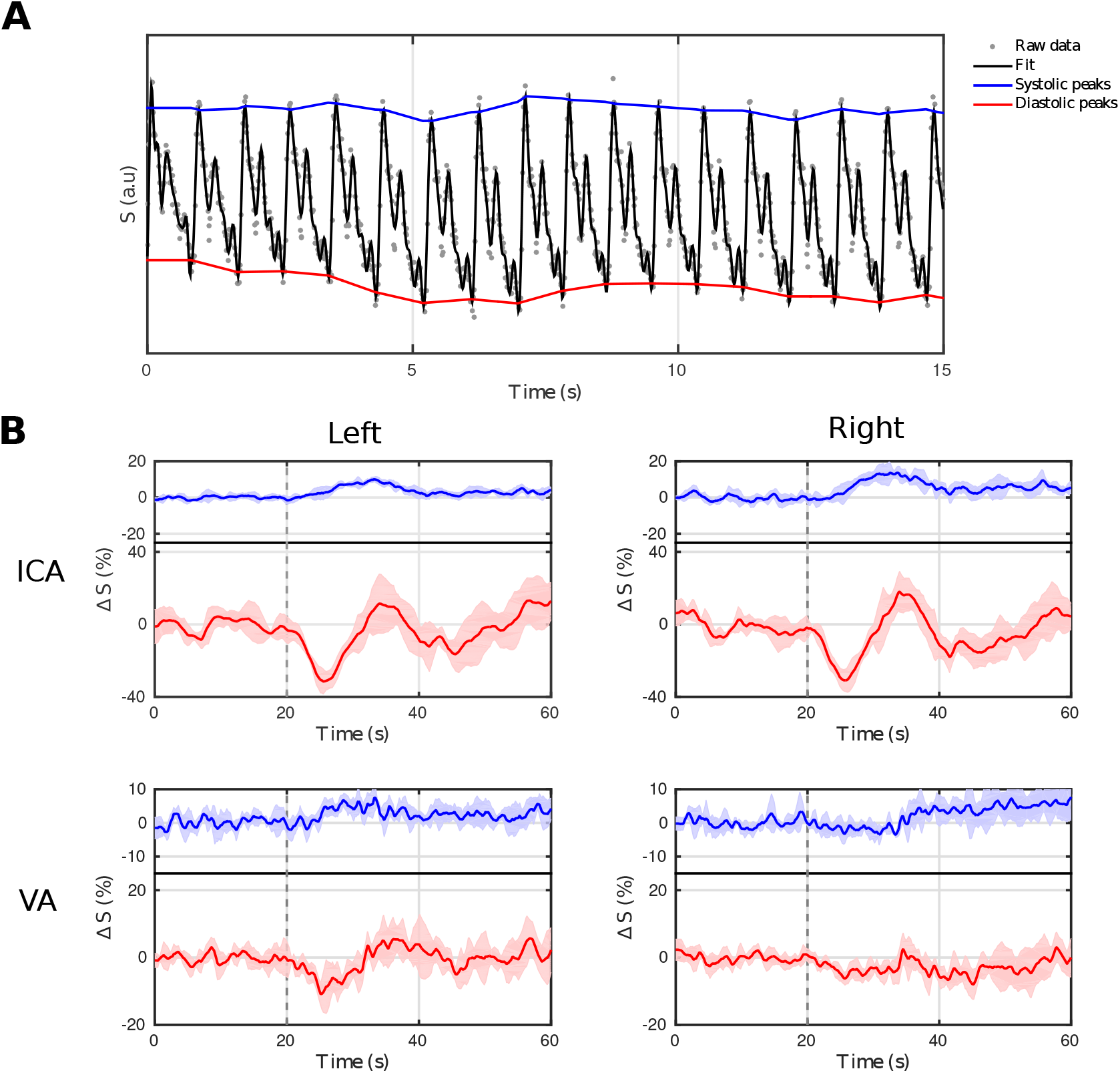
A) First 15s of the *TCR-challenge* time series taken from the left ICA ROI. B) The *systolic* (blue) and *diastolic* (red) peak time series (i.e. signal envelope) averaged across the five repeats, for both ICA and VA bilaterally. The onset of the thigh-cuff release is highlighted with a dotted grey line. Both left and right ICA show clear time locked responses to the *TCR-challenge* in both *systolic* and *diastolic* time series. The same responses are far less evident in the VA time series, particularly in the right side.

## 4. Discussion

This study outlines a new approach for real-time measurement of arterial flow, which we have dubbed Dynamic Inflow MAgnitude Contrast (DIMAC). The inflow effect causes the magnitude of the complex GRE MR image to be inherently sensitive to CBFV, and this sensitivity scales inversely with TR. Thus, the use of very short TRs, which are a prerequisite for high temporal resolution imaging that can resolve the pulsatile component of flow, has the fortuitous consequence of also imparting high sensitivity to CBFV in the signal. So when combined with a high flip angle, short TR GRE imaging enters the DIMAC regime of sensitivity, which we have proposed is a valuable source of image contrast for measuring dynamic pulsatile arterial flow. Using a two-compartment model to simulate the spoiled GRE signal, we have shown that in the DIMAC regime the signal shows high relative sensitivity to pulsatile CBFV with respect to pulsatile CBV. Bianciardi et al. previously delineated three different regimes of image contrast with respect to sensitivity to temporal fluctuations in CBFV (Bianciardi et al., 2016), and focussed specifically on the regime in which there is minimal sensitivity to CBFV, and thus the dynamic image contrast predominately reflects pulsatile changes in CBV. In contrast, in this study we have considered the regime with sensitivity to pulsatile CBFV. Our simulations show that with very short TRs and high flip angle we can achieve great sensitivity to changes in CBFV, with minimal contribution from CBV, even in instances where there is a significant extraluminal partial volume present. By limiting ourselves to single slices that are carefully positioned perpendicularly to large cerebral arteries, we have shown that in the DIMAC regime we can measure arterial flow with high temporal resolution in real-time.

### 4.1. Dynamic measurements

A major advantage of the DIMAC approach we have outlined is the ability to measure pulsatile flow in real-time, and resolve individual beat-to-beat pulsatile waveforms. This is desirable as averaging across multiple cardiac cycles, as is common with PC-MRI CBFV waveforms, typically leads to significant morphological differences compared with TCD (Wagshul et al., 2011), best characterised as a loss of fine structure, i.e. fewer clearly resolved distinct peaks and troughs. These key features (peaks and troughs and their relative timings) that have been used to characterise the CBFV waveform in the TCD literature (Aggarwal et al., 2008; Kurji et al., 2006; Lockhart et al., 2006; Robertson et al., 2008), are often integral to attempts to indirectly derive higher order measures of the cerebrovascular system such as intracranial pressure and cerebral perfusion pressure (Aggarwal et al., 2008), or downstream compliance (Robertson et al., 2008). To counter this problem, recent technological advances have been exploited in an increased effort to develop PC-MRI acquisitions with sufficiently high temporal resolution to enable real-time pulsatile flow measurement (Markl et al., 2016). These novel real-time acquisitions have mostly focussed on cardiac applications (Joseph et al., 2012; Klein et al., 2001; Nayak et al., 2000; Nezafat et al., 2005), but there are still some neuroscience applications, in particular focussing on CSF dynamics (Chen et al., 2015; Yildiz et al., 2017). As these real-time acquisitions allow dynamic flow variations and beat-to-beat variability to be assessed during short segments of time, the respiratory cycle’s effect on flow can be easily deduced (Chen et al., 2015), and a more complete assessment of pulsatile flow can be made. For the same reason real-time approaches are crucial for accurately gauging pulsatile flow in the presence of atrial fibrillation (Markl et al., 2016), and quantifying the effect of beat-to-beat variations on flow characteristics. A a benefit of DIMAC over PC-MRI is that it negates the need for additional gradients to be played out, and so more readily facilitates high temporal resolution acquisitions. Real-time PC-MRI methods using EPI readouts typically achieve temporal resolutions between 50-80ms (Chen et al., 2015; Yildiz et al., 2017). This situation can be improved slightly with non-Cartesian approaches (~40 – 50 ms) (Joseph et al., 2012; Kowallick et al., 2014), but it is still not comparable with the temporal resolution of 15ms used here, and there is potential for further improvements if non-Cartesian approaches are also developed for DIMAC. The benefit to higher temporal resolution is that it allows the higher order harmonics of the pulsatile component of the signal to be accurately sampled, and high fidelity pulsatile waveforms to be more accurately measured and in a shorter time, even on a single beat basis. In particular, the precision with which the relative timings can be measured between different peaks and troughs in the flow waveform is fundamentally dependent on temporal resolution, and it is well established that relative timing information within the pulsatile flow waveform contains physiologically valuable information (Asgari et al., 2019; Naqvi et al., 2013).

Real-time acquisitions are necessary to study flow responses to dynamic physiological challenges, such as Valsalva and Mueller manoeuvres (Kowallick et al., 2014; Thavendiranathan et al., 2012), allowing pulsatile flow to be assessed in the context of naturalistic physiological stress. Such approaches are beneficial for cerebral flow applications, as determining the mechanisms of autoregulation is highly active research area (van Beek et al., 2008). Furthermore, low frequency modulation of arterial tone appears to be especially prevalent in the brain, and the existence of slowly evolving fluctuations in CBF are well established cross-species, and have been recorded using numerous modalities (Obrig et al., 2000). Understanding how these low frequency changes modulate beat-to-beat pulsatile dynamics is a potentially exciting new research direction that real-time pulsatility methods like DIMAC offer. This point is highlighted by the *TCR-challenge* experiment presented in this study, in which we measure the induced flow response to an orthostatic challenge, and of note is the fact that the temporal dynamics of the *systolic* and *diastolic* peak time series show differential responses. Such flow dynamics can only be determined by methods with sufficient temporal resolution to accurately resolve the pulsatile component of flow.

### 4.2. Flow velocity

The basis of DIMAC signal contrast is high sensitivity to flow velocity. The *v_c_* parameter sets the upper limit of CBFV sensitivity and is determined by TR and slice thickness, parameters which are themselves limited by both practical and hardware considerations. This is a non-trivial limitation of the approach described, as in contrast with PC-MRI, in which the comparable velocity encoding (VENC) parameter can be set relatively independently from the other image acquisition parameters, DIMAC sensitivity is directly determined by those parameters. One inherent limitation with DIMAC when measuring pulsatility, is that fluctuations in CBFV that exceed the critical velocity vc do not translate into signal variance, i.e. sensitivity to flow velocity saturates. This signal saturation results in a total loss of velocity information above *v_c_,* unlike PC-MRI, where misspecification of the VENC parameter leads to aliasing that can in some cases be corrected (Xiang, 1995), albeit with an SNR cost. The parameters used here predict a *v_c_* of ~ 67 cm s^-1^, which is below the peak velocity one might expect to measure for ICAs, and on the limit for MCAs (Brant, 2001), which risks losing sensitivity to the most central lamina of the cross-sectional flow distribution. This is also a current limitation when comparing pulsatile flow waveforms across different individuals, or physiological conditions such as with the experiment presented here, particularly in the case of the *HC-challenge*. If during systole a fraction of spins in the central lamina exceed *v_c_*, then the observed differences between hypercapnia levels may be more attributable to a graded response of this effect. This effect results in a non-linear DIMAC signal behaviour in high flow velocity situations, and so future research will be needed to better understand this and the degree to which it confounds pulsatility estimates. However, in instances of slower flow velocities this becomes less of an issue, and so perhaps there is more potential in using this method to target smaller arteries.

A useful attribute of PC-MRI is that with careful consideration of acquisition and analysis protocols, flow velocity can be quantified in meaningful units. This has clear benefits, particularly from a clinical perspective. In principle a quantitative estimate of CBFV could be extracted from the DIMAC contrast as we have demonstrated in the *Supplementary material* with a simple flow phantom experiment. Equation 1 can be solved for velocity if an estimate of the M_0_ of arterial blood is provided, which could be achieved using a separately acquired image with a long TR such that the critical velocity v_c_ is sufficiently small so that the majority of flowing blood spins exceed it. The main barrier to this would be limited spatial resolution and associated extra-luminal partial volumes, which lead to multiple sources of signal contrast that present a challenge for quantifying pulsatility in physiologically relevant terms (Viessmann et al., 2017). However, as demonstrated by the simulations presented in Fig.1, in the DIMAC regime the signal from static spins is effectively nulled, due to the saturating effect of a train of short interval, high flip angle RF pulses, suggesting static tissue partial volumes only contribute marginally to the total signal variance. However, this topic warrants further focussed investigation in order to fully delineate the effects of different signal sources, and thus explore whether CBFV might be quantified with an acceptable level of precision.

### 4.3. Target arteries

In this feasibility study, we have focussed on large intracranial arteries, however there may also be value in focussing DIMAC on smaller vessel applications. The need to isolate the pure arterial blood signal phase makes partial volume errors non-negligible in PC-MRI (Nayak et al., 2015), whereas inflow related techniques are less impeded by extra-luminal signal contributions given that the contrast itself depends on them being relatively saturated. Our simulations suggest that while the total signal magnitude is inversely correlated with the extra-luminal partial volume, the relative sensitivity to pulsatile CBFV over CBV changes is relatively unaffected. Furthermore, the cerebral large arteries are relatively stiff in comparison to the aorta (Mitchell, 2008), which also contributes to this effect as consequently they show only relatively small changes volume as a function of cardiac pulsations. Moving down the cerebral arterial tree the relative stiffness is also expected to increase (Hayashi et al., 1980), which further reduces the impact of CBV changes. Thus, the relative insensitivity to extra-luminal partial volumes in the DIMAC signal, at least in terms of pulsatile CBFV sensitivity, and the inherently small pulsatile CBV changes in cerebral arteries, both implies that the method could be optimised to measure pulsatility in smaller arteries with sub-voxel diameters. A smaller artery application would also benefit from ultra-high field, not only due to the general increase in SNR, but also the specific beneficial effect on inflow contrast, which is now well established in the TOF angiography literature (Grochowski and Staśkiewicz, 2017).

## 5. Conclusion

The pulsatile nature of arterial blood flow provides a clinically relevant insight into arterial structure/function and its effect on cerebral health. In this study, we demonstrate the feasibility of a new approach to measuring pulsatile flow in cerebral arteries by exploiting the inflow effect that is present in highly accelerated GRE acquisitions. In addition to presenting simulation results to support this new method, we also present in-vivo data from major cerebral arteries (ICA, VA and MCA), but suggest that this technique may be used to target smaller vessels that are beyond the scope of other methods. Our hypercapnia challenge results provide evidence that the DIMAC signal is sensitive to subtle changes in vascular tone, and we have shown that this method allows beat-to-beat assessment of pulsatile flow without requiring averaging across cardiac cycles. Furthermore, using a thigh cuff release challenge we have demonstrated that this real-time approach allows the full range of dynamics, including both pulsatile and non-pulsatile components, associated with transient flow responses to be measured. Thus, we believe this novel DIMAC method provides a promising new approach for studying cerebral arterial function, which will ultimately be valuable in researching arterial function in ageing and cerebrovascular disorders.

## 6. Acknowledgments

This work was funded in whole, or in part, by the Wellcome Trust [WT200804]. For the purposes of Open Access, the author has applied a CC BY public copyright license to any Author Accepted Manuscript version arising from this submission. Thanks to Peter Weale for valuable input during pilot scanning.

