## Supplementary material for "Measuring arterial pulsatility with Dynamic Inflow MAgnitude Contrast (DIMAC)": SupplementaryMaterial_MRM_final.pdf

### 1. Supplementary material

#### 1.1 Flow phantom experiment

The flow dependence of DIMAC was validated in an experiment using a custom-made flow phantom. The phantom consisted of a plastic tub serving as a reservoir, a plastic tube (0.6mm inner diameter), a DC motor driven high-pressure water pump powered by a 24V power supply and controlled with a motor controller, a flow meter, and a Raspberry Pi unit used to program the motor controller. The pump, motor controller, Raspberry Pi, and reservoir were all placed outside the magnet room, with the tubing fed through the waveguide to create a circuit. Inside the magnet room the tubing was wound into a coil that was placed outside the head coil with the end forming a straight length that is encased in a 3D-printed agar filled container inside the head coil. A 40/60 glycerin/water mix was used to ensure a flow medium with similar viscosity to blood, which is expected to result in a laminar flow profile within the straight section of the tubing within the head coil.
The experiment consisted of adjusting the flow rate through the phantom by varying the voltage supplying the water pump, starting at 100% of maximum, down to 15% in increments of 5%, giving 18 discrete flow velocities. After each voltage adjustment the flow was allowed to stabilize and then data were acquired from a single transverse DIMAC slice, positioned at isocentre perpendicular to the tubing inside the phantom, for approximately 15 s (1024 repeats). For each repetition an additional scan was obtained for the purposes of estimating the $M_0$ value of the flow medium, with the same parameters as the default protocol but with a TR=2 s and only 2 repetitions.

##### 1.1.1 Flow phantom experiment analysis

Raw data were reconstructed using custom in-house scripts in MATLAB, including phase correction, GRAPPA reconstruction and coil combination using the square root of the sum of squares. A single voxel contained within the tubing of the phantom was identified by choosing the brightest voxel in the mean image across all repetitions. A quantitative estimate of the flow velocity ($v$) was obtained by scaling the region of interest (ROI) voxel signal intensity from the flow sensitive scan ($M_z$)

with the signal intensity from the $M_0$ scan by rearranging equation 1. The flow medium $T_1$ was assumed to be 1.1 s (Ohno et al., 2017) although given that TR << $T_1$ this has a very small effect on the flow velocity estimate within the plausible range of $T_1$ values.

DIMAC estimates of $v$ ($v_{DIMAC}$) were compared with flow meter estimates ($v_{meter}$) using least-squares linear regression, and the coefficient of determination ($R^2$) was calculated to determine the degree of shared variance between the two flow velocity estimate methods. A Bland-Altman plot was used to judge the level of agreement, with the mean difference reflecting any systematic bias. A one-sample t-test on the difference between $v_{meter}$ and $v_{DIMAC}$ was used to test for systematic differences. For all statistical tests, data points where $v_{meter} > v_c$ were excluded, as the theory predicts that $v_{DIMAC}$ values plateau for values > $v_c$, i.e. flow sensitivity is lost and thus $v_{meter}$ and $v_{DIMAC}$ values will not agree.

#### *1.1.2 Flow phantom experiment results*

Fig.3 shows the results of the flow phantom experiment, which demonstrates the strong flow velocity dependence of the DIMAC signal, as it increases monotonically with flow velocity as expected. A linear fit between the two estimates results in an $R^2$ of 0.94, indicating an extremely high degree of shared variance. However, a high level of shared variance does not necessarily indicate good agreement between two different estimates of the same property. A Bland-Altman plot is often used to make a judgment on the level of agreement, as shown in in Fig.2. There is a small yet significant average bias of -5.19 cm $s^{-1}$ ($p<0.001$) between estimates ($v_{meter} – v_{DIMAC}$), with 95% limits of agreement of ±10.78 cm $s^{-1}$.

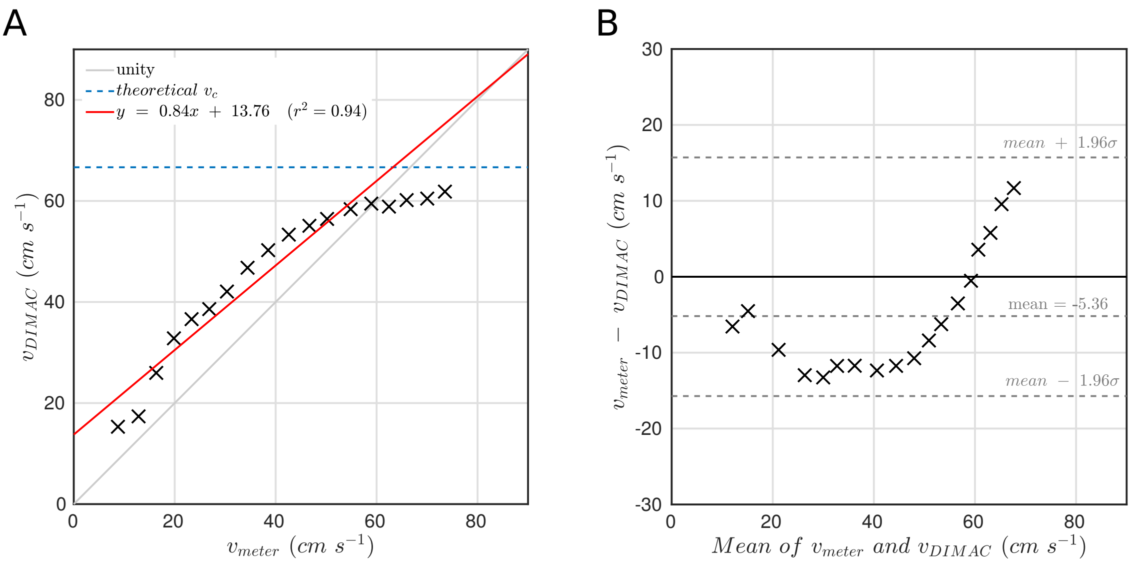


**S1: A) DIMAC estimated flow velocity estimates ($v_{DIMAC}$) plotted as a function of flow meter estimates ($v_{METER}$) with line of best fit. Y-error bars are not included for the DIMAC estimates for the purpose of clear presentation, as the variance in the measurements is very small and so the error bars are not visible on the chosen scale. B) Bland-Altman plot showing 95% limits of agreement between $v_{DIMAC}$ and $v_{METER}$.**

#### *1.1.4 Flow phantom experiment discussion*

Using a simple flow phantom we have demonstrated a monotonic relationship between flow velocity and the DIMAC signal as expected. Furthermore, we show how the signal can be scaled to give reasonable quantitative flow velocity estimates. However, it is clear that there is a small systematic bias that leads to moderate disagreement between the measurements, and the DIMAC values plateau lower than the theoretically predicted value. However, this is perhaps not particularly surprising given the relatively rudimentary nature of the experiment and the numerous sources of error in both measurements. The experiment was performed primarily for demonstration purposes and as a simple validation of the theory, but the results do suggest that with further research efforts, quantitative blood flow velocity estimates may be made precisely with the DIMAC method.

### *1.2 HC-challenge experimental timings*

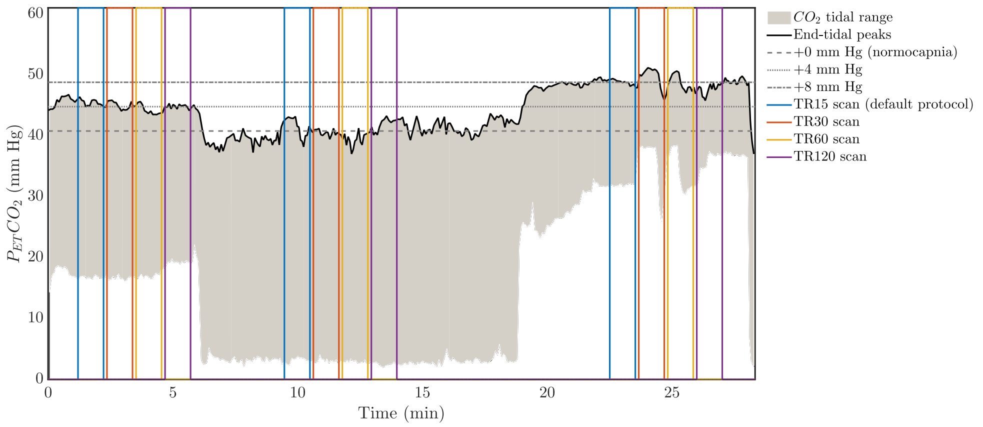


**S2: Example subject $P_{ET}CO_2$ trace. For this subject the HC conditions were in the order of HC4, HC0, HC8. It can be seen that when transitioning to a new HC level, sufficient time was given to allow a new steady-state to be reached, which ensured that each HC level was distinct from every other, with only small fluctuations about the target $P_{ET}CO_2$ level. As expected, fluctuations around the HC0 (normocpania) level are of a slightly greatly magnitude, as this is the only level which can't be explicitly controlled through gas delivery (i.e. the subjects just breath medical air).**

### *1.3 TCR-challenge physiological response*

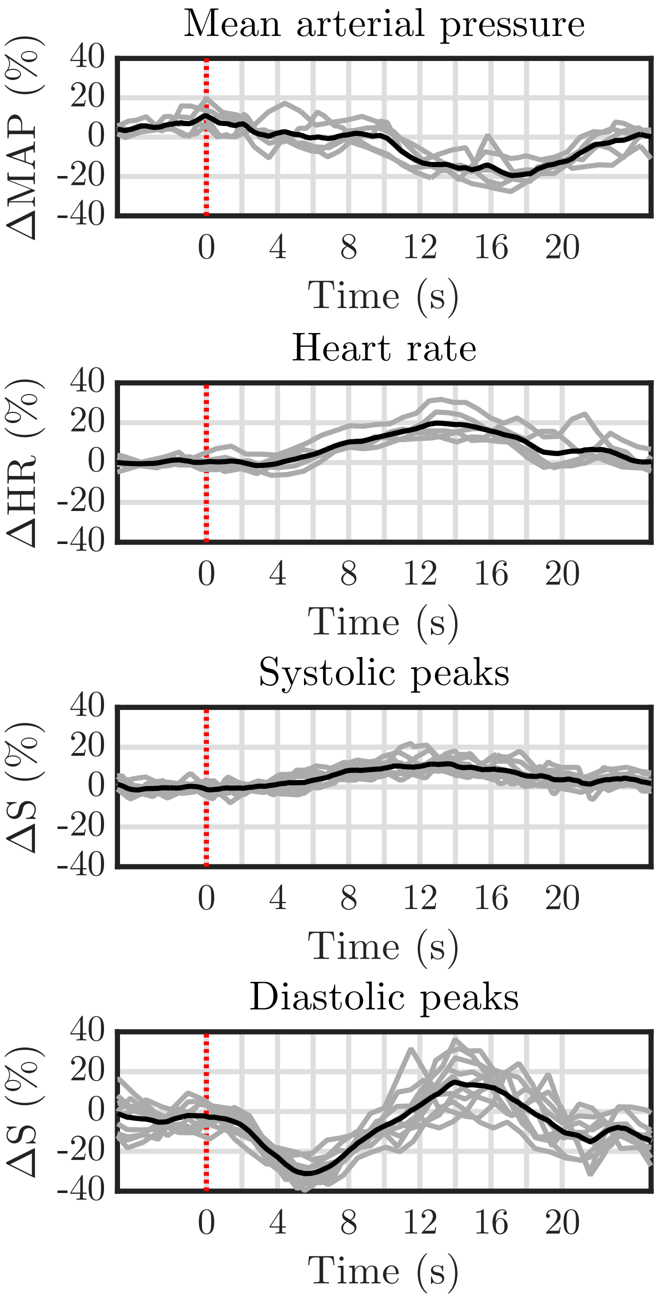


**S3: Time locked physiological (MAP and HR) and ICA *systolic* and *diastolic* peak responses to the TCR-challenge. The dotted red line indicates the onset of the TCR. For each panel, the 5 grey curves are the individual repeats the black curve is the average across repeats. The first (top) panel shows that there is TCR evoked reduction in mean arterial pressure (MAP) that peaks with ~ 20% reduction at ~16s post stimulus. The second panel shows that there is a TCR evoked increase in heart-rate (HR) that peaks with ~20% increase at ~14s post stimulus. The third panel shows that there is a small sustained rise and fall in the *systolic* peaks time series, which peaks at ~10% increase at ~14s. The fourth (bottom) panel shows that there is a pronounced decrease in the *diastolic* peaks time series, which peaks ~30% reduction at ~5-6s, followed by a return to baseline and subsequent overshoot, which peaks ~10-15% increase at ~14s.**
